# Social feedback and the emergence of rank in animal society

**DOI:** 10.1101/018374

**Authors:** Elizabeth A. Hobson, Simon DeDeo

## Abstract

Dominance hierarchies are group-level properties that emerge from the aggressions of individuals. Although individuals can gain critical benefits from their position in a hierarchy, we do not understand how real-world hierarchies form, or what signals and decision-rules individuals use to construct and maintain them in the absence of simple cues. A study of aggression in two groups of captive monk parakeets (*Myiopsitta monachus*) found a transition to large-scale ordered aggression occurred in newly-formed groups after one week, with individuals thereafter preferring to direct aggression against those nearby in rank. We describe two mechanisms by which individuals may determine rank order: inference based on overall levels of aggression, or on subsets of the aggression network. Both pathways were predictive of individual decisions to aggress. Based on these results, we present a new theory, of a feedback loop between knowledge of rank and consequent behavior, that explains the transition to strategic aggression, and the formation and persistence of dominance hierarchies in groups capable of both social memory and social inference.

## Introduction

Individuals from social species must interact with each other to reproduce, find food, and survive. Higher-level social structures such as hierarchies emerge when interacting individuals need to manage trade-offs in the costs and benefits of social associations [1, 2]. One of the most important is the dominance hierarchy, where group-wide “global” rankings are derived from local aggressive interactions and form emergent social properties [3–5].

Aggression has obvious immediate costs, including energy expended and the possibility of injury. Benefits, conversely, can be both immediate and delayed. Individuals may fight to gain immediate access to contested resources, or they may aggress in order to gain rank, which then provides these individuals with delayed rank-dependent benefits. Most importantly, aggression that results in higher dominance rank often increases an individual’s access to foraging resources and reproductive opportunities (*e.g.*, Ref. [6–8]).

Groups across a broad range of taxa are structured by dominance rank despite large variations in cognitive skills. Dominance hierarchies are found in primates [9, 10], social carnivores [11, 12], ungulates [13, 14], birds [15–18], fish [19], and even crustaceans [20, 21]. These group-level social structures form and stabilize on the basis of perceptions and actions necessarily made at the individual level [22]. Dominance rank is generally achieved through a series of aggressive events, and hierarchy formation takes place without top-down control in a manner that is largely independent of intrinsic properties of the individuals involved [23–25].

Previous experimental and theoretical work on dominance hierarchies shows that interaction outcomes can shape how individuals interact in subsequent interactions [26], that individuals may use a jigsaw approach to determine how to interact with a few individuals based on observations of event outcomes [23, 27], and that observed win/loss outcomes can be strong determinants of an individual’s choice of targets of future aggression [28–30].

Recent evidence suggests that individuals in many species have the cognitive ability to use their observations of others to inform their own behaviors, including several primate species [9, 10, 31–33], ravens [34], hyenas [35], and fish [36–39]. However, while this evidence shows that individuals observe each other and react to these observations, we do not currently understand how individuals integrate information on the outcomes of their own interactions with observations of others’ interactions in order to determine rank.

We are thus faced with two distinct questions. How does rank become relevant to individual decisions to aggress? And what cues might individuals use in order to learn emergent social properties like rank?

This paper presents three findings, based on detailed, highly-resolved observations of aggression in a social avian species. In doing so, we provide an account of the interaction between local decision-making and global properties. Our first finding is a strong signal of the influence of rank on decision-making. This is seen in how aggression is allocated: target choice became structured over and above what is necessary to reproduce the rank structure alone. This occurred around a week after initial group formation. After this transition, individuals preferentially directed aggression towards those nearby in rank and avoided interactions with those far below them in rank.

Our second finding is that in this structured society, both levels of aggression and subsets of the full network (network motifs in the form of aggression chains) provided cognitively-accessible signals of rank. These pathways are the likely mechanisms through which rank is inferred. Rank is a global property, but in these structured systems can be learned by judicious observation of local interactions. Our third finding is that the motif pathway not only provided a signal of relative rank, but was strongly predictive of actual behavior. Individuals were far less likely to direct aggression against the terminal individual in an observable aggression chain.

Taken together, these results help explain the emergence of rank as an interaction between two processes: inference of rank from cognitively-accessible social signals, and decision-making that correlates with these signals. They indicate a critical role played by a knowledge-behavior feedback loop—between inference of group-level properties, and consequent decision-making. Such feedbacks may be a critical pathway for how evolved systems reduce uncertainty by tying together multiple timescales [40]; our findings here have parallels in discoveries of how signal use in primates tracks coarse-grained features of a social network [22, 41].

These results provide new insight into the problem of choosing targets and establishing a dominance hierarchy in species that that lack simple perceptual cues, such as size or spatial location, for an individual’s rank. In these more complex societies, rank order is necessarily a cognitive construct that summarizes the many dyadic-level interactions into an emergent group-level property.

Our results derive from studies of two independent groups of monk parakeets (*Myiopsitta monachus*), a small neotropical parrot native to temperate South America and notable for its highly social colonial and communal nesting behavior [42] as well as its widespread success as an invasive species [43–46]. Monk parakeets exhibit several characteristics of complex sociality [47, 48], and to our knowledge, is the first parrot species in which detailed and quantitative dominance hierarchy analysis has been conducted [47].

Parrot species represent an intriguing comparison to dominance hierarchy studies conducted in primates and humans. They share many characteristics with primates, such as large relative brain size and advanced cognition [49, 50], extended developmental period [49, 51], long lifespans [52–54], and individualized relationships within complex social groups [47, 55]. Additional characteristics, such as vocal learning and high fission-fusion dynamics, are uncommon in most primates [56], but are shared by parrots and humans [57]. As such, understanding the ways in which parrots form and maintain dominance relationships in complex social groups has the potential to further our understanding of rank in socially and cognitively complex species.

## Methods

### Data Collection Protocols

Our study is based on observation of directed aggression in groups of monk parakeets housed in captivity at the U. S. Department of Agriculture National Wildlife Research Center in Gainesville, Florida. We formed two independent groups (*N* = 21 and 19) and observed aggressive events during novel group formation (additional details in Refs. [47, 48]). Prior to our study, parakeets were housed in groups of one to six individuals per cage; while some were in visual contact, direct physical contact between individuals in different cages was not possible. To facilitate individual identification, we marked each bird with a unique facial pattern using colored nontoxic permanent markers (Sharpie, Inc.). We collected morphometric data to determine whether rank could be predicted by an individual’s size.

Each captive group was released sequentially into a 2025 m^2^ semi-natural outdoor flight pen and observed over the course of 24 days by 1-4 observers. We used all occurrence sampling [58] to record data on directed agonistic behaviors. As in Ref. [47], we restricted our analysis to dyadic events with clear outcomes, where an aggressor physically displaced or supplanted a target individual. This resulted in a “win” for the aggressor and a “loss” for the target of the aggression. We also collected nearest-neighbor data [47] to determine how rank affected spatial association patterns.

We divided the 24-day study period into four 6-day study quarters to facilitate comparisons across the two replicate social groups.

### Estimation of dominance hierarchies

We use Eigenvector Centrality (EC, Ref. [59]) on the directed aggression networks as our primary means of determining rank. EC considers both the direct and indirect links in aggression networks in a recursive fashion to determine individual power: high centrality equates to low power. Individuals have lower power if they are the target of aggression or if they are the target of aggression from another individual with low power, and so forth. EC is one of the main algorithms for determining consensus beliefs within a network [60] and dominance ranks based on EC power scores are strongly correlated with the main alternative ranking methods such as I&SI [61]. EC allows for the direct quantification of power, rather than just a linear order; this means it can distinguish cases where two individuals are “nearly equivalent” from those where differences in dominance are reliable and large.

EC is closely related to David’s Score (DS; Ref. [62]); both have found widespread uses in the characterization of dominance relations. In the terminology of [63], EC and DS are “depth” measures of network consensus, weighting the interactions between two individuals *i* and *j* in ways that depend on interactions each individual has had with others. An alternative are “breadth” measures that predicate the rank or power of an individual *i* based only the interactions that *i* engages in. When measuring the effect of signal consensus (a related, but distinct, problem to that of establishing measures of power from aggression data) Ref. [63] found EC comparable to the breadth measure Weighted Simple Consensus (WSC), both of which performed well. In our data, EC correlates strongly with all three measures; see SI Text.

### Average Rank Aggression

We are most interested in how individuals make decisions on how to direct aggression based on relative rank. To measure this, we use average rank aggression, *R*(Δ). *R*(Δ) is the average number of aggressive acts, per day, directed at an individual Δ ranks away, and is defined as:

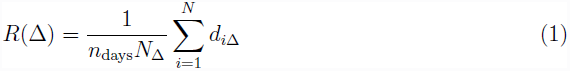

where *d*_*i*Δ_ names the individual whose rank is Δ steps from *i*’s rank, *N*_Δ_, the total number of individuals who have a potential target Δ ranks away, and *n*_days_ the total number of days used to construct the **d** matrix. The Δ can be positive or negative, and indicates an individual of lower or higher rank than *i* respectively.

Average rank aggression is our primary signal of individual decision-making. We are interested in determining whether the observed rank aggression indicates the influence of rank on decision-making. In order to do this, we construct null models for the range of behaviors we expect to see if individuals interact in a unstructured fashion.

In a group of *n* individuals, there are *n*(*n* − 1) free parameters in an aggression network, but only *n* − 1 numbers are required to specify that network’s EC power scores. EC (indeed, any ranking or scoring system) thus amounts to a lossy compression of the original data, summarizing the behavioral patterns relevant to the establishment of a dominance hierarchy. Conversely, for any given dominance hierarchy there are many possible behavioral patterns. Our null model for aggression is defined as a random sample from this larger set; in addition, we preserve the total aggression of each individual (see SI Text).

Any particular sample from this null will preserve (on average) the EC power scores, but will be otherwise unstructured and contain no correlations that are unnecessary to preserve those scores. We can measure relevant properties, such as *R*(Δ), on these null networks. Deviations in the real data from that null indicate individuals are systematically directing aggression in ways that differ from what is otherwise expected for an aggression network with that dominance structure.

### Signals of rank

Estimating rank via EC is computationally challenging. As noted by Ref. [63], breadth measures such as WSC are of particular interest because they correlate with these more sophisticated measures, but are more likely to be cognitively-accessible to individuals within the system. For this reason, we also measure WSC on the directed aggression network. The WSC score of an individual is the product of the total amount of aggression (number of events) received, multiplied by the number of distinct individuals who directed that aggression. We order individuals by these dominance scores to determine individual rank. As in the case of EC, a higher WSC equates to lower rank because these individuals receive more aggression from more individuals.

Individuals may use breadth-based signals such as WSC, but they may also use measures sensitive to other features of the network. In particular, even though EC rank is a property of the aggression network as a whole, small portions of that network may contain signals of relative rank, and these smaller network subsets may more easily perceived.

To study information of this second kind, we focus on a particular kind of motif—the aggression chain—where we can trace a line of aggression from individual *i*, to individual *j*, to individual *k*, and so forth. We measure the signals contained in such chains using average weighted rank difference, *W*(*n*). *W*(*n*) quantifies the extent to which an aggression chain provides information about relative rank difference for chain length *n*. *W*(2) is defined as

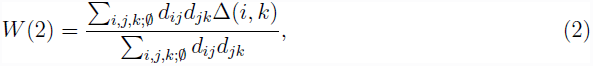

where Δ(*i*, *k*) is the rank difference between *i* and *k*, and 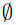 indicates that in any instance, the identities of individuals *i*, *j*, and *k* do not overlap. *W*(2) is then the average rank difference between any two individuals, weighted by the product of aggression seen along all chains connecting them.

*W*(2) takes into account not only the existence of the chain, but also its strength. In general, weighted network ties are generally more informative about the social relationships among individuals and are more robust to sampling differences [48, 64–66] than relying solely on presence-absence information. *W*(*n*) for *n* larger than two is defined similarly (see SI Text); the number of possible motifs we need to examine to compute *W*(*n*) grows exponentially with depth, and we stop our analysis at *n* equal to six. (Note that while the total number of motifs grow exponential with *n*, *W*(*n*) quantifies the average amount of information in any particular aggression chain, not the total amount of information in all chains.)

When *W*(*n*) is significantly different from the null, this indicates the *presence* of information in aggression chains over and above what is expected from systems with the same power scores, but otherwise unstructured aggression.

### Use of Motif-based signals of rank

To determine if individuals actually perceived and used these motifs as a signal of rank, we measure the behavioral signatures of transitive inference. Transitive inference occurs when one individual uses knowledge about its own interactions with a target (*j*) and third-party observations of how *j* interacts with additional targets (*k*, *l*, *m*,…) to infer its own likelihood of winning over one of *j*’s targets, such as *k*. For the case of three individuals, *i* and *k* would have a transitive relationship with individual *j* if the amount of aggression directed from *i* to *j* (*d*_*ij*_) and the aggression from *j* to *k* (*d*_*jk*_) was related to the amount of aggression *i* directed to *k* (*d*_*ik*_).

We tested for transitive relationships between individuals in the first and last positions anchoring each aggression chain (Fig. 1), up to a chain length of 6. We quantified fractional transitivity, *T*(*n*), for a chain length of two as:

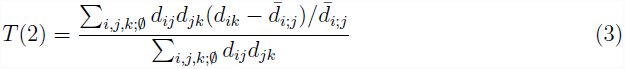

**Figure 1:**
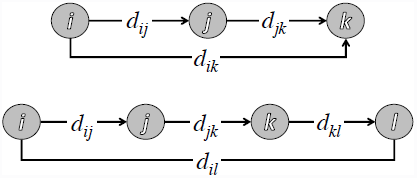
Aggression chains of length two and three; *W*(2) quantifies the extent to which aggression along the chain (*d*_*ij*_ and *d*_*jk*_) is a signal of rank difference between *i* and *k*; *T*(2) quantifies the extent to which *d*_*ik*_ is higher, or lower, than average aggression by *i*, given aggression levels *d*_*ij*_ and *d*_*jk*_.

where 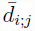 is the average aggression *i* directed towards all individuals other than *j*, and 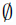 indicates that our sums exclude cases where the identities of individuals *i*, *j*, and *k* overlap. The natural extension to *n*-step chains, *T*(*n*), is defined in the SI Text. *T*(*n*) is positive or negative when an individual *i* increases or decreases, respectively, its aggression against *k* given that *k* is at the end of an *n*-step chain.

Very positive *T*(*n*) means that individuals prefer to aggress against those at the end of chains while very negative *T*(*n*) means that individuals preferentially avoid such aggression. When *T*(*n*) is significantly different from null, this indicates the *use* of information in aggression chains over and above what is expected from systems with the same power scores, but otherwise unstructured aggression.

### Data exclusions

Our initial analysis indicated that differences in aggression patterns in Group One and Group Two were strongly driven by a single individual, NBB. This individual was persistent in her attempts to affiliate with others in Group Two, but was not able to form a strong affiliative relationship within the group [47]. Aggression directed at NBB appeared to be less due to a strategic choice of target and more a reaction to NBB’s persistent and apparently unwanted attempts to affiliate. We conducted a post-hoc analysis of aggression patterns and found that excluding this single individual from Group Two resulted in a pattern of aggression that was much more consistent with that observed in Group One. We present the results for Group Two including NBB in the SI to show how unusual decision-making by this individual affects the overall patterns created by the other 18 individuals.

## Results

We recorded a total of 1013 aggressive events in Group One and 1360 events in Group Two. Despite the captive conditions, some individuals avoided interacting which each other, resulting in incompletely connected aggression networks in both groups (density of 0.89 in Group One; 0.92 in Group Two). We find no evidence that rank could be reliably determined based on simple underlying rules such as size or spatial proximity. Rank was not significantly associated with the physical size of individuals, including weight, wing length, and beak properties (|*r*^2^| < 0.18, *p* > 0.05, SI Figure 1). In addition, spatial proximity was a poor signal of relative rank (|*r*^2^| < 0.12, SI Figure 2).

**Figure 2:**
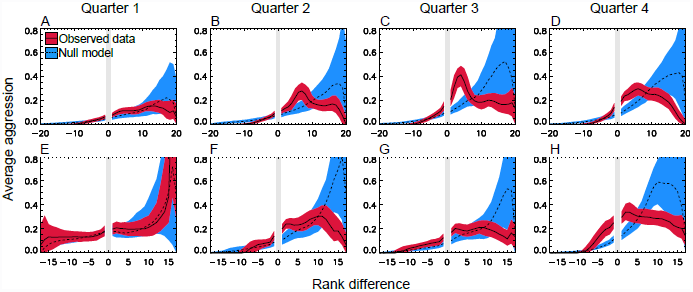
The emergence of structured aggression. Average rank aggression (*R*(Δ),Eq. 1) for Group One (a-d) and Group Two (e-h) across study quarters; each quarter is six days long. Patterns of aggression (red band/solid line) in both groups are consistent with the null (dashed line/blue band) in the first Quarter, but diverge in Quarters two, three and four as individuals focus their aggression on those nearby in rank. Points are averaged ±1 rank; bands are 1σ bootstrap-estimated error ranges. Maximum rank difference in Group Two (*N* = 19) is 17 because one individual was dropped. See SI Text.

### Emergence of social structure

Average rank aggression in both Groups One and Two was initially consistent with the null model during the first quarter of the study period (Fig. 2a and 2e). However, behaviors quickly diverged from null expectations as individuals began to structure their behavior in ways strongly correlated with rank. Aggression patterns in the final three-quarters of the study period all diverged strongly from null expectations (Fig. 2b–d and 2f–h), with aggression strongest towards individuals nearby in rank. We refer to this as rank-focused aggression. Combining the final three quarters of the data increases this signal (Fig. 3a and 3b).

**Figure 3:**
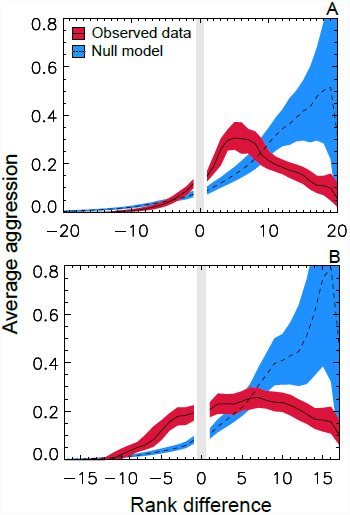
Structured aggression. Average rank aggression (*R*(Δ), Eq. 1) pooled over the final 3 quarters of the study period for Group One (a) and Group Two (b). Pooling data provides a gain of signal-to-noise that further refines our understanding of the strategies at work once the behavior-knowledge feedback is in place. Both groups show a focusing of aggression towards those nearby in rank, over and above null expectation. Higher levels of rank entrepreneurship—aggression directed, contrary to expectation, upwards in the hierarchy— are visible in Group Two. Points are averaged ±1 rank; bands are 1σ bootstrap-estimated error ranges.

This is our first main result: the onset of strong deviations from the nulls points to the emergence of structured behavior at the individual level dictated, at least in part, by relative EC rank.

### Inferring Rank: The Magnitude Pathway

Determination of EC is computationally intensive. In our data, EC correlated strongly with WSC (*r*^2^ ≈ 0.73 in both groups) and thus knowledge of WSC can provide at least partial knowledge of EC. Because of WSC’s reliance on levels of aggression alone, rather than network structure, we refer to this mechanism as the “magnitude pathway”.

We look for signals of the use of this pathway, by looking for evidence that aggression is structured as a function of relative WSC rank. We find evidence for reduced aggression at individuals widely separated in WSC rank (large positive Δ) compared to the null. This indicates that, in additional to providing knowledge of rank, WSC-derived rank is also predictive of some features of individual aggression. However, we do not see strong evidence for increased aggression to those nearby in WSC rank—none in Group One, and only weak evidence in Group Two (see SI Text).

The fact that WSC was predictive of aggression indicates that the magnitude pathway may play an important a role in structuring aggression. However, the absence of a signal of rank focused aggression implies that there were patterns of aggression *invisible* to the breadth-based WSC measure. If individuals used WSC signals to help direct their aggression to those nearby in rank, they must have been supplementing them with other sources of information. The absence of rank-focused aggression in the WSC case is an example of how rank defined by reference to WSC does not capture all of the structure relevant to individual aggression.

### Inferring Rank: The Motif Pathway

Weighted rank difference, *W*(*n*), allows us to investigate whether social information was encoded within chains of aggression (Fig. 4). We refer to this as the “motif pathway”.

**Figure 4:**
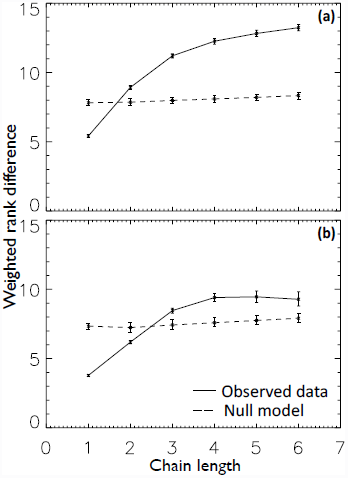
Cognitively-accessible social properties. Shown is the weighted average rank difference for different aggression chain lengths (*W*(*n*), Eq. 2) for Group One (a) and Group Two (b). Solid line shows the observed signal, while the dashed line shows that found in the null, with 1*σ* error ranges. Significant evidence exists for a strong signal of relative rank contained in aggression chains. Such signals are then accessible to individuals capable of transitive inference.

In the null (dashed line in Fig. 4), chain length encodes little or no information about relative rank. Individuals tend to be lower-ranked than their aggressors, but seeing an individual at the end of a long chain provides little or no additional information about its rank. By contrast, the observed data (solid line in Fig. 4) encoded significant information in aggression chains. In both Groups One and Two, local motifs contained a substantial amount of global information.

An observation of an individual at the end of a longer chain (length ≥ 3) provided more information about relative rank than an observation of an individual at the end of a short chain (length 1 or 2). This encoding potentially allowed individuals to distinguish between individual nearby (Δ ∼ 5) and distant (Δ > 10) in rank—a discrimination impossible in the nulls.

This is our second main result: network motifs, in addition to breadth-based magnitude measures, can provide signals of relative rank.

Fractional transitivity, *T*(*n*), allows us to investigate whether these aggression chains are predictive of actual behavior. Our analysis found a strong difference between the null model and the observed data (Fig. 5). In the null (dashed line), chains predict increased aggression: if A aggresses against B, and B against C, this leads A, on average, to direct increased aggression against C. This is independent of chain length—both long and short chains predict similar levels of increased aggression.

**Figure 5:**
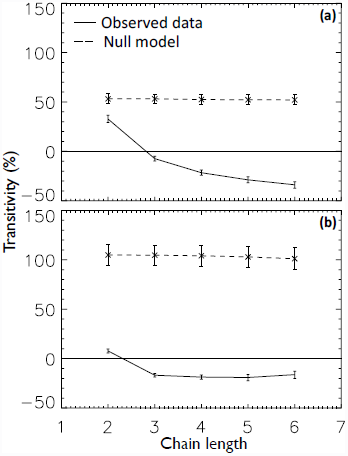
Aggression chains predict behavioral preferences. In both Group One (a) and Group Two (b), aggression against those at the end of an chain was reduced compared to null. For aggression chains three steps or longer, we find preference inversion: a negative transitivity meaning that (on average) A directed less-than-average aggression against D if D was found at the end of a chain A→B→C→D.

In contrast, behavior in the observed data (solid line) showed the opposite effect. While short chains predicted a (small amount) of increased aggression (*T*(2) greater than zero), long chains were associated with reduced aggression (*T*(*n*) less than zero for *n* greater than two). This remarkable inversion is what is expected when individuals use the information content of motifs to predict relative rank, and then both (1) preferentially avoid conflict with much lower-ranked individuals and (2) focus aggression on rank neighbours.

This is our third main result: network motifs predict behavior.

### Quarter-by-quarter Dynamics of *W*(*n*) and *T*(*n*)

Having demonstrated the existence and predictive power of signals, we can also ask about the time-frame over which the signals themselves emerge by looking at *W*(*n*) quarter-by-quarter. In Group One we can already find evidence for the existence of these signals, over and above null expectation, at the one-step and two-step level, in the first quarter. The signals were present, but *R*(Δ) results suggest that they are not yet used by participants. By contrast, the signal is almost entirely absent in the first quarter for Group Two. After the first week (*i.e*., for the later quarters) the signal became significantly stronger, covered a wider range of ranks, and we saw signals at the three-step, and often at the four-step, level in both groups (see SI Text).

We can also examine *T*(*n*) quarter-by-quarter to look for the dynamical emergence of behavioral preference. The first quarter showed some evidence for the use of motif information to structure behavior, with shifts in preferences of on average 32 percentage points relative to the null in both groups. Effects became larger at later times, with behavioral preferences shifting by on average 82 percentage points relative to the null in the final three quarters. Preference inversion appears only in these later quarters. The emergence of this behavioral pattern parallels the emergence, over time, of the large scale order seen in Fig. 2, and of cognitively-accessible signals seen in Fig. 4 (see SI Text).

## Discussion

Our results show how group social structure forms, how cognitively-accessible signals allow for inference of rank, and how these signals predict core features of actual behavior. Our study provides insight into two longstanding questions about dominance in complex groups: how does dominance emerge, and how do individuals infer and act upon it?

We found that (1) a transition towards more structured aggression occurred rapidly, about a week after initial group formation, (2) behavioral motifs and chains of aggression provided information about relative rank, and (3) symbolic distance along aggression chains was predictive of aggression preferences. We found that use of the magnitude pathway, WSC, could only explain some of this structure, but that individuals could supplement this with depth-based information in the form of aggression chains.

The patterns seen in Fig. 3 are thus driven, on the one hand, by information about rank (in both the magnitude pathway, and the motif pathway seen in Fig. 4), and, on the other hand, by decisions to aggress that we track in Fig. 5. We have found that both the motif signal, and the extent to which it predicts aggression patterns, increase over time, paralleling the emergence of the large-scale order seen in Fig. 3.

Our results imply the existence of a feedback between behavior and knowledge. Aggression at the individual level leads to large-scale facts about dominance rank at the group level. Individuals are able to gain knowledge about these ranks, and to use this information to adjust their behavior accordingly.

This leads to the diachronic emergence of global social signals, including the dominance ranks that have been a central part of study in animal behavior. This feedback loop is dependent on individuals possessing two critical cognitive skills: social memory and social inference. Individuals must be able to identify individuals and remember the outcomes of past events, and then integrate these memories to structure subsequent behavior. This feedback loop may play a key role in dominance hierarchy development in larger, more complex social groups.

We focused on two distinct knowledge pathways: (1) the magnitude pathway, which requires observers to track incoming aggression to each individual, regardless of source identity and (2) the motif pathway, which requires partial knowledge of aggression network structure.

Under the magnitude pathway, the number of aggressors and the total amount of aggression targeted at each individual is tracked to determine rank. As more interactions occur, more information exists upon which rank can be inferred. Although the magnitude pathway is more likely to be cognitively accessible to individuals [63], decisions made on the basis of this pathway alone are unable to explain all of the behavior we see. Depth-based measures provide additional discriminatory power, and are predictive of actual aggression including the rank-focused aggression seen in Fig. 3.

Once these signals are in place, effects are strong. Aggression levels differed by a factor of two or more from the null model (Fig. 3), signals allowed individuals to discriminate across nearly the full range of relative ranks (Fig. 4), and these signals predict shifts in behavioral preferences of 50 percentage points or more (Fig. 5).

Significant features of the social signal are contained in the two-and three-step aggression chains. That observed shifts in behavior (quantified by *T*(*n*)) can be due to direct reliance on the signal (detected by *W*(*n*)) is supported by experimental studies. These find that larger symbolic distances are not only salient, but actually more easily perceived than neighboring pairs [67,68]. Meanwhile, our detection of rank focused aggression is consistent with results that find nearby ranks perceptually salient, including the finding that rank reversals occurring at small relative rank differences are perceived as more stressful [34].

Spatial assortment may be a signal of rank in some systems [69]. However, we found no evidence that parakeets use either spatial assortment or physical size to structure their aggression. Instead, our work suggests that the formation of a hierarchy relies on cognitive inference over complex relationships. Our work has found that aggression motifs contain a substantial amount of information that could be used infer relative rank.

Despite being more cognitively demanding, the motif pathway explains structuring of aggression over and above that from the magnitude pathway. Use of this signal requires that individuals reason inductively about relationships over and above observed pairwise comparisons, an ability known as transitive inference. While transitive inference (A beats B, B beats C, therefore A beats C) is more sophisticated than recall of pairwise comparisons (A beats B), experimental work has established that a wide range of social species have the ability to infer or learn indirectly [70]. Evidence for transitive inference has been documented in both non-social [67, 71, 72] and social contexts [73–75]. Meanwhile, observational studies find evidence for the use of higher-order (*i.e*., beyond pairwise) strategies where actions of one individual against another are influenced by third parties, both in individual and group-level decision-making and perception [37,38,76–78]. Reliance on transitive motifs provides a natural source of such rules: a decision to aggress against a target can be influenced by the aggression that others display against the target.

Previous work has shown how relatively simple rules can be used to govern dyadic or triadic interactions at a local scale. Here, we have considered how hierarchies form on larger scales in more complex social groups. As group size increases, the number of relationships that must be tracked to determine group-level rankings increases dramatically. Our results here help explain how this complexity can be managed by use of cognitively-accessible pathways, and how the use of such pathways may feed back to how individuals choose to aggress.

The question of cognitive complexity is crucial. A human observer equipped with sophisticated computational tools can determine rank order given knowledge of all pairwise aggressive events. However, it is highly unlikely that the individuals have the cognitive ability needed to use these methods, and they must rely in part on simpler, “ecological” [79–81] methods such as the magnitude and motif pathways described above.

The ability to construct a model of the dominance hierarchy, based on a combination of direct experience and indirect observation of others, could be particularly adaptive in species where dispersal results in the regular integration of new members into existing groups and in groups with high fission-fusion dynamics. Reliance on, and consequent amplification of, cognitively-accessible signals means that individuals would not have to directly interact with all possible combinations of individuals in the group.

Indirect inference of dominance rank can allow individuals to predict the behavior of others while conserving energy and reducing the possibility of injury [82]. It can also facilitate integration of immigrants into existing groups, and allow more rapid formation of dominance hierarchies. This may be particularly adaptive in species where dispersal results in the regular integration of new members into existing groups, and in groups with high fission-fusion dynamics.

## Conclusions

How individuals come to know their social worlds, and how that knowledge feeds back to influence social properties, is a crucial part of group dynamics. This paper has tracked the emergence of strategically directed aggression, the signals that could enable it, and how these signals predict decisions to aggress.

Previous work on transitive inference by non-human animals has often focused on experimental manipulation of trained subjects [67, 70]. These experiments are generally removed from social situations. They can provide critical evidence that individuals have the necessary cognitive skills, but usually cannot show a direct link between those abilities and an understanding of the social landscape of an actual group and its effect in real-world situations. Our work provides new evidence for the importance of transitive inference in the real-world problem of directing aggression.

Our findings on the information contained in chains of aggression and the strategic use of this information allow us to construct a mechanistic account of both what signals of rank in observed behavior might be available to individuals (the knowledge pathway) and how these signals influence decision-making (the behavior pathway). It allows us to explain the dynamical transition as the onset of a complex interaction between knowledge of rank and consequent behavior, and a closing of a knowledge-behavior feedback loop that leads to the formation of complex society.

## Author summary

An individual’s success depends critically on socially-constructed properties such as rank. A detailed study of two independent captive parakeet groups reveals how these properties come into being. We show that individuals can use localized patterns in the aggression network to learn the relative ranks of individuals, and that these signals of rank strongly correlate with individual decisions to aggress. Over time, feedback between knowledge and behavior leads to the emergence of strategic aggression: individuals focus their aggression on those nearby in rank.

## Acknowledgments

We thank Dr. Michael Avery and the entire staff of the Florida Field Station for their help and support for this research, especially Kandy Keacher’s help with animal husbandry, and our field assistants D. John, T. McIntosh, and A. Hobson for data collection and enthusiasm. S.D. thanks Kirstin G.G. Harriger for helpful conversations. E.A.H. was supported by the New Mexico Higher Education Graduate Fellowship, Loustaunau Fellowship, and National Science Foundation (NSF) GK-12 DISSECT (#DGE-0947465) Fellowship, research grants from the Associated Students of New Mexico State University, American Ornithologists’ Union, Sigma Xi, and the NMSU Biology Graduate Student Organization, with further support from NSF Grant #IOS-0725032 and associated REU supplement to Timothy Wright. Part of this work was conducted while E.A.H. was a Postdoctoral Fellow at the National Institute for Mathematical and Biological Synthesis, an Institute sponsored by the NSF, the U.S. Department of Homeland Security, and the U.S. Department of Agriculture through NSF Award #DBI-1300426, with additional support from the University of Tennessee, Knoxville. S.D. and E.A.H. were supported in part by NSF Grant #EF-1137929, and by the Santa Fe Institute. All animal activities conducted during this study were approved by the New Mexico State University Institutional Animal Care and Use Committee (protocol number 2006-027).

## Supplementary Material for “Social feedback and the emergence of rank in animal society”

### 1 Definitions of *W*(*n*)and *T*(*n*)

As in the main text, we defined the weighted rank aggression as

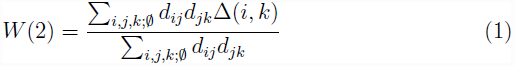

where we write 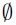 to indicate the restriction to distinct values of *i*, *j*, and *k*. Higher orders can be defined in a similar fashion, so that

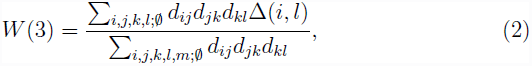

and

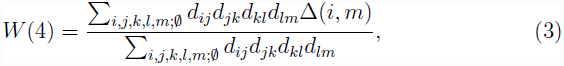

and so forth, always restricting to non overlapping choices for the indices.

**Table 1:** Correlations of power scores (in the case of David’s Score and WSC) and overall rank (in the case of I&SI, which provides only an ordering, not a score). In all cases, correlations are significant at *p* < 0.001. Analysis for final three quarters of the data.

|  | Group One | Group Two | Type |
| --- | --- | --- | --- |
| EC to David's Score | 0.96 | 0.95 | Pearson $r^2$ |
| EC to WSC | 0.73 | 0.73 | Pearson $r^2$ |
| EC to I&SI | 0.93 | 0.76 | Spearman |

Similarly, the transitivity is defined as

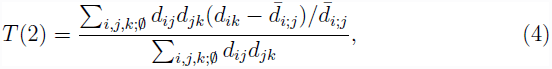

where 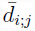 is the average aggression directed by *i* against individuals other than *j*. We exclude *j* from this average so as not to induce a spurious correlation. and higher orders can be defined as

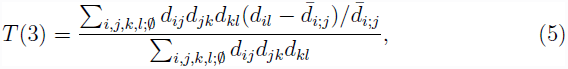

and

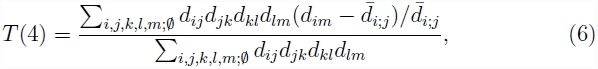

and so forth, always restricting to non overlapping choices for the indices.

### 2 Correlations between Eigenvector Centrality and other measures

Eigenvector Centrality (EC) is our main way to characterize the dominance hierarchies of both our systems. It correlates strongly with other measures of rank, as shown in Table 1, where we compare EC to David’s Score, I&SI, and Weighted Simple Consensus.

**Figure 1:**
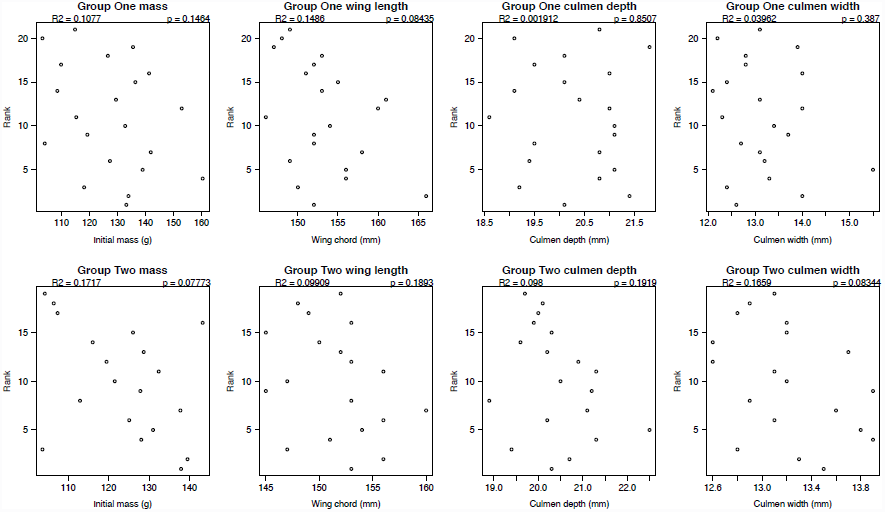
Eight null results. Physical characteristics do not signal rank.

### 3 Physical Size and Rank: Null Results

We looked for correlations between four main physical cues—mass, wing length, culmen depth and culmen width—and rank. In all four cases, and in both groups, we found no evidence of any rank signal. Correlations were close to zero, and consistent with the null, each case. See Fig. 1. (Culmen measurements refer to the size of the beak; depth and width are both measured at the nares, which is roughly the base of the beak.)

### 4 Spatial Proximity and Rank: Null Results

If birds nearby in rank tend to spend more time physically near each other, spatial proximity could serve as a signal of rank. In particular, we would expect a negative correlation between (absolute) relative rank difference between *i* and *j*, and the number of times *i* was observed to be *j*’s nearest neighbour in space. We find, however, that spatial proximity is at best a poor signal of relative rank. Even in the final three-quarters, where behavior is most regular, in Group One, proximity provides only a weakly anticorrelated signal (*r*^2^ = −0.14); in Group Two, it provides no signal at all (*r*^2^ = 0.04). If we exclude pairs who allopreen, the correlations become weaker still: group one has *r*^2^ = −0.12; Group Two has *r*^2^ = −0.02. This is shown in Fig. 2.

**Figure 2:**
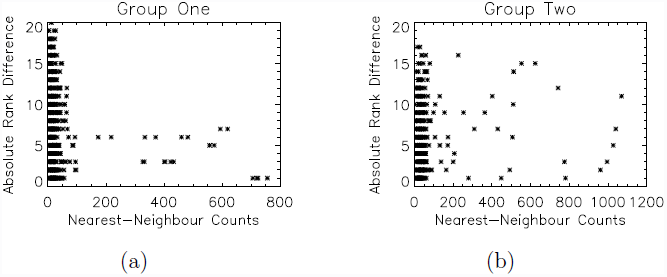
Two null results. Spatial proximity is not a strong signal of rank in either Group One (*r*^2^ = −0.12) or Group Two (*r*^2^ = −0.02). Note that each point here represents a pair; there are thus 210 points in Group One, and 171 points in Group Two, on these two plots, with the vast majority of points overlapping near zero on the *x* axis.

### 5 Null Models for Aggression

To compute power, EC maps the pattern of directed aggression to a Markov process. We define the matrix **d**, where an element *d*_*ij*_ is the number of times individual *i* aggressed against *j*. From **d**, we can calculate the transition matrix **t**

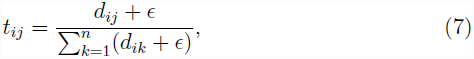

where *t*_*ij*_ is the probability that if *i* is observed aggressing, it is aggressing against *j*, and *ϵ* is a small regularizing term. The power distribution, *v*, is the stationary distribution of this transition matrix; equivalently, the dominant left-eigenvector of the matrix **t**.

There are *n*(*n* – 1) free parameters in an aggression network, but only *n –* 1 integers are required to define a linear ranking system. Thus, any ranking system amounts to a lossy compression of the original data, summarizing the behavioral patterns relevant to the establishment of a dominance hierarchy. Conversely, for any given dominance hierarchy there are many possible behavioral patterns. Our null model for aggression is defined as sampling randomly from this set.

In particular, because there are many possible **d** and **t** matrices compatible with a particular power distribution *v*, we can define a null model as random draws from the set of matrices that have, on average, the same *v*. Now we constrain not only the linear ranking, but also a power-score.

We construct a measure over this set implicitly, sampling from by means of the following algorithm.

1. Take *k* is equal to 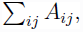 *i.e*., the total number of observed aggressions in the data.
2. Construct a *k*-element time-series, *T*, of individuals, σ_1_,σ_2_…σ_*k*_, where the σ_*i*_ are drawn by means of iid samples from *E*_1_(**d**).
3. Re-order *T* so that it contains no consecutive pairs, *i.e*., so that σ_*i*_ ≠ σ_*i*+1_.
4. Use the Expectation-Maximization (EM) algorithm [2] to find the transition matrix **t***'* for a Markov Model consistent with this time-series. Since *t*_*ii*_ is equal to zero, our reordering will fix *t*_*ii*_ equal to zero and the final solution **t***'* found by the EM algorithm respects this constraint.
5. Convert **t***'* to an aggression series **d***'* by fixing 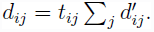.

The time-series {*σ*_*i*_} itself can be interpreted as a linked chain of observed aggression pairs, σ_*i*_ → σ_*i*+1_, σ_*i*+1_ → σ_*i*+2_, and so forth, but this time series should not be interpreted as a potential observation; correlations *between* aggressive acts in the series are not constrained to those observed in the real world.

A particular null transition matrix **t***'* represents aggression preferences, which we then map to a set of expected behaviors, **d***'*, or a set of counts that could have been observed in the wild. We assume the same overall aggression levels for each individual seen in the original data *i.e.*, 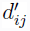 is equal to 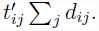. We generated 10,000 null replicates.

In our figures reporting *R*(Δ) we bin neighbouring ranks (*i.e.*, we estimate *R*(Δ) by averaging the raw measurements for Δ − 1, Δ and Δ + 1, reflects RMS uncertainties in the ranks themselves; when Δ is equal to one, we average 1 and 2, and similarly for Δ equal to negative one. This improves signal to noise in each bin at the cost of correlating bins. To determine errors in the observed data (red bands in our Figures 2 and 3 in the main paper), we use the statistical bootstrap and resample with replacement from the original observed data, reporting median and 1σ bands. As in the case of the null model, we bin neighbouring ranks.

### 6 Group Two with NBB

As described above, we removed NBB from our analyses of Group Two; here, for completeness, we show the late-time stationary properties of Group Two including NBB in Fig. 4. Immediately apparent is a peak in the average aggression directed towards much lower ranks.

Multi-modal observations allow us to attribute these effects entirely to NBB’s anomalous attempts to affiliate with high-ranking individuals who continually repelled these unwanted attempts. Exclusion of this single individual completely eliminates this peak, as can be seen in Fig. 3 in the Main Text. Inclusion of NBB also leads to an apparent delay in the onset of structured aggression in Group Two.

### 7 Weighted Simple Consensus

Ref. [1] described a simple, cognitively accessible measure of network consensus called Weighted Simple Consensus. Under WSC, an individual’s score is the product of the total amount of aggression (number of displacements) received, multiplied by the number of distinct individuals who directed that aggression. WSC power correlates strongly with EC power in both groups (*r*^2^=0.73 for the final three quarters, in both groups; *r*^2^=0.70 quarter-by-quarter average), and because of its relative simplicity may thus play the role of a signal of relative rank—in particular, individuals may use WSC to estimate relative rank, and use those estimates to guide their decision-making. If WSC does indeed play the role of a rank indicator that individuals use to direct aggression, we should see an imprint of this decision-making in the data itself; in particular, we should see that average aggression is focused, over and above the null, as a function of WSC rank. Indeed, this is what is found in both groups, as shown in Fig. 5: we find strong, above null signals that relative rank, when estimated by WSC, predicts focused aggression, most obviously in the suppression of long-range rank aggression.

**Figure 3:**
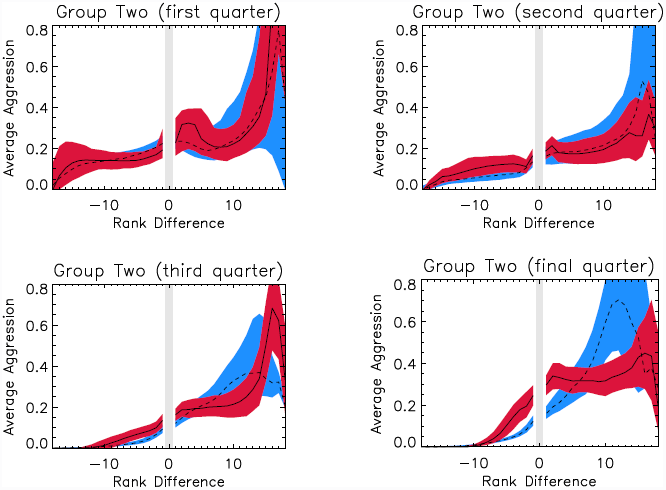
Including NBB in the analysis of Group Two leads to an apparent delay in the onset of structured aggression as aggression against NBB drives it down the hierarchy.

**Figure 4:**
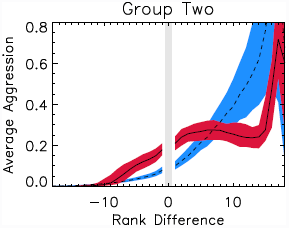
Stationary features of Group Two are largely unchanged by the inclusion of NBB, except for the peak at the largest rank differences corresponding to aggression directed against the lowest ranking individual, NBB, by high-ranking individuals who repel NBB’s unwanted attempts at affiliation.

**Figure 5:**
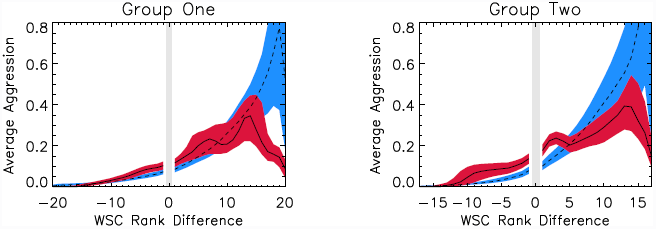
Weighted Simple Consensus as a guide to focused aggression in Group One (top panel) and Group Two (bottom panel); final three quarters. Red band is average aggression as a function of rank difference (and 1*σ* errors); Blue is the EC null as before.

Comparing Fig. 5 and Fig. 3 of the main paper, we see that, while (1) WSC is a predictor of EC rank, and (2) WSC is correlated with rank-focused behavior, it is also the case that rank focusing is much stronger under the EC measure than under WSC. In group one, for example, rank aggression directed at rank separations less than five is a factor of 1.95 above null when using EC ranks, but only 1.34 above null when using WSC ranks; meanwhile, in both group one and group two, there is greater suppression of long-range rank aggression when using EC ranks compared to WSC ranks.

Given this greater focusing, if individuals were using WSC directly, it would require unusual fine-tuning for them to pattern their usage to achieve a tighter focus in EC than they do in WSC itself. This leads us to consider the possibility, in the main paper, that individuals may also rely on depth-based measures of relative rank.

### 8 Quarter-by-quarter Dynamics of *W*(*n*) and *T*(*n*)

The main text shows the emergence of large-scale strategic aggression structures (Fig. 2, Main Text), as well as the existence of local signals (Fig. 4, Main Text) and behavioral preferences (Fig. 5, Main Text) that could be driving this shift. In this section, we examine the quarter-by-quarter dynamics of the local signals.

Figs. 6 and 7 show that, to a limited extent, some of these local signals are already available within the first week. In Group One, Quarter One, for example, two-step chains provide additional information about relative rank over and above one-step chains; note, however, that additional steps in the chain do not significantly improve knowledge of relative rank over observation of a two-step chain. In Group Two, signals in Quarter One are extremely weak, with the difference between a one-step and a five-step chain of less than three ranks, and with no statistically significant gain from going from one-step to two-step chains.

**Figure 6:**
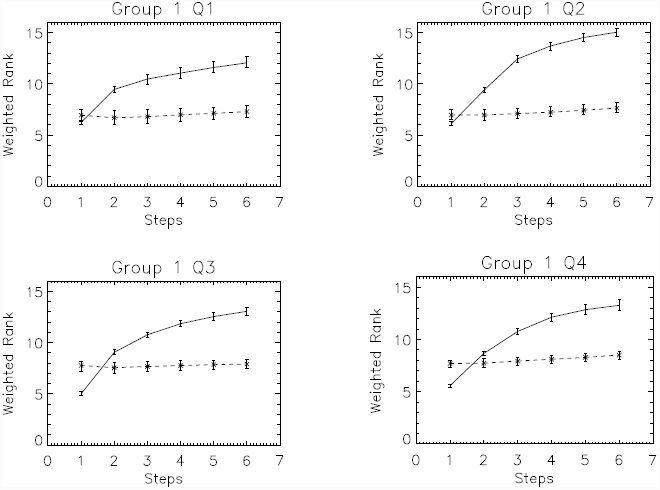
*W*(*n*) broken out quarter-by-quarter for Group One. Solid line: observed data; dashed line: null. Transitive signals of rank already appear in Quarter One, where the one-and two-step aggression chains can clearly distinguish relative ranks. In Quarters Two, Three and Four, these signals strengthen, so that three- and even four-step aggression chains now contain additional information.

In both cases, signals are much stronger, and remain much stronger, after the first week, with Quarters Two, Three, and Four in both groups having significant information of relative rank in the one-, two- and three-step chains, and (to a more limited extent) in the four-step chains.

**Figure 7:**
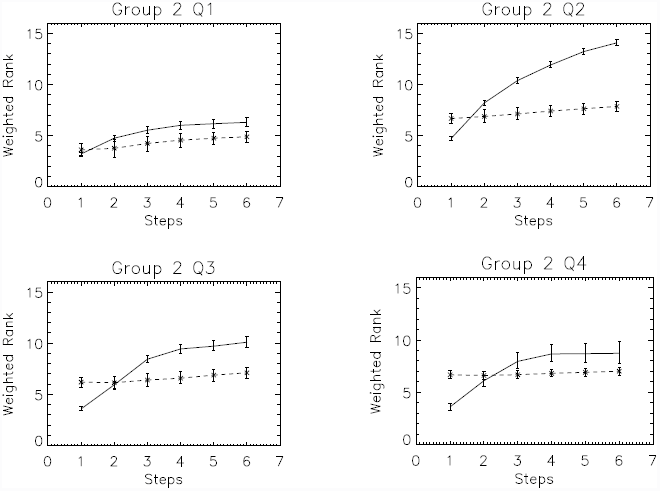
*W*(*n*) broken out quarter-by-quarter for Group Two. Solid line: observed data; dashed line: null. Very little information exists in Quarter One; In Quarters Two, Three and Four, chains up to length three and even four (in Quarter Two) contain distinct information.

**Figure 8:**
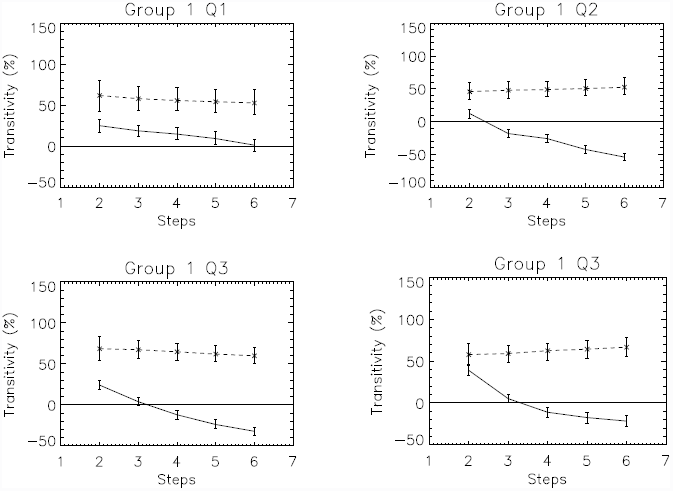
*T*(*n*) broken out quarter-by-quarter for Group One. Solid line: observed data; dashed line: null.

**Figure 9:**
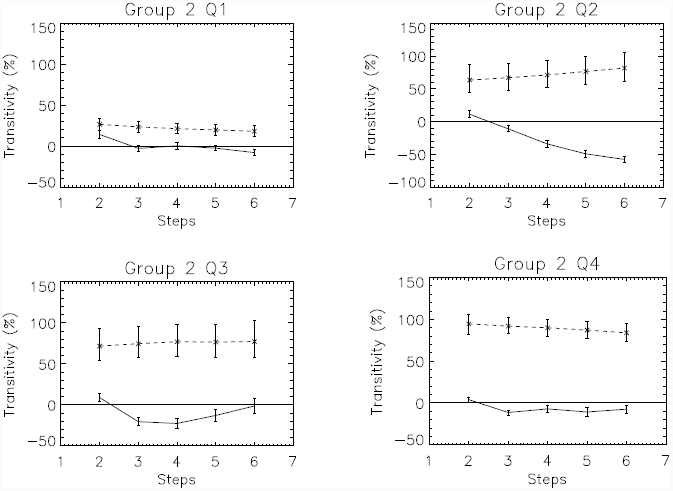
*T*(*n*) broken out quarter-by-quarter for Group Two. Solid line: observed data; dashed line: null.

